# Bacteria are everywhere, even in your COI marker gene data!

**DOI:** 10.1101/2021.07.10.451903

**Authors:** Haris Zafeiropoulos, Laura Gargan, Sanni Hintikka, Christina Pavloudi, Jens Carlsson

**Affiliations:** Department of Biology, University of Crete, Voutes University Campus, Heraklion, Greece; Hellenic Centre for Marine Research (HCMR), Institute of Marine Biology, Biotechnology and Aquaculture (IMBBC), Heraklion, Crete, Greece; Area52 Research Group, School of Biology and Environmental Science/Earth Institute, University College Dublin, Dublin 4, Ireland

**Keywords:** environmental DNA (eDNA), metabarcoding, software tool, tree of life (tol), unassigned sequences, container, Docker

## Abstract

The mitochondrial cytochrome C oxidase subunit I gene (COI) is commonly used in eDNA metabarcoding studies, especially for assessing metazoan diversity. Yet, a great number of COI operational taxonomic units or/and amplicon sequence variants are retrieved from such studies and referred to as “dark matter”, and do not get a taxonomic assignment with a reference sequence. For a thorough investigation of this dark matter, we have developed the Dark mAtteR iNvestigator (DARN) software tool. A reference COI-oriented phylogenetic tree was built from 1,240 consensus sequences covering all the three domains of life, with more than 80% of those representing eukaryotic taxa. With respect to eukaryotes, consensus sequences at the family level were constructed from 183,330 retrieved from the Midori reference 2 database. Similarly, sequences from 559 bacterial genera and 41 archaeal were retrieved from the BOLD database. DARN makes use of the phylogenetic tree to investigate and quantify pre-processed sequences of amplicon samples to provide both a tabular and a graphical overview of phylogenetic assignments. To evaluate DARN, both environmental and bulk metabarcoding samples from different aquatic environments using various primer sets were analysed. We demonstrate that a large proportion of non-target prokaryotic organisms such as bacteria and archaea are also amplified in eDNA samples and we suggest bacterial COI sequences to be included in the reference databases used for the taxonomy assignment to allow for further analyses of dark matter. DARN source code is available on GitHub at https://github.com/hariszaf/darn and you may find it as a Docker at https://hub.docker.com/r/hariszaf/darn.

**Author summary:** DARN is a software approach aiming to provide further insight in the COI amplicon data coming from environmental samples. Building a COI-oriented reference phylogeny tree is a challenging task especially considering the small number of microbial curated COI sequences deposited in reference databases; e.g ~4,000 bacterial and ~150 archaeal in BOLD. Apparently, as more and more such sequences are collated, the DARN approach improves. To provide a more interactive way of communicating both our approach and our results, we strongly suggest the reader to visit this [Google Collab notebook](https://colab.research.google.com/drive/1XorHsBm1uqx5TTZsH7SeVRkUA2SS8dnY?usp=sharing) where all steps are described step by step and also this GitHub page where our results are demonstrated. Our approach corroborates the known presence of microbial sequences in COI environmental sequencing samples and highlights the need for curated bacterial and archaeal COI sequences and their integration into reference databases (i.e. Midori, BOLD, etc). We argue that DARN will benefit researchers as a quality control tool for their sequenced samples in terms of distinguishing eukaryotic from non-eukaryotic OTUs/ASVs, but also in terms of understanding the unknown unknowns.

## Introduction

DNA metabarcoding is a rapidly evolving method that is being more frequently employed in a range of fields, such as biodiversity, biomonitoring, molecular ecology and others (Deiner et al. 2017, Ruppert et al. 2019). Environmental DNA (eDNA) metabarcoding, targeting DNA directly isolated from environmental samples (e.g water, soil or sediment, Taberlet et al. 2012a), is considered a holistic approach (Stat et al. 2017) in terms of biodiversity assessment, providing high detection capacity. At the same time, it allows wide scale rapid bio-assessment (Stat et al. 2017) at a relatively low cost as compared to traditional biodiversity survey methods (Ji et al. 2013).

The underpinning idea of the method is to take advantage of genetic markers, i.e. marker loci, using primers anchored in conserved regions. These universal markers should have enough sequence variability to allow distinction among related taxa and be flanked by conserved regions allowing for universal or semi-universal primer design (Deagle et al. 2014). In the case of eukaryotes, the target is most commonly mitochondrial due to higher copy numbers than nuclear DNA and the potential for species level identification. Furthermore, mitochondria are universally present in eukaryotic organisms, and can be easily sequenced and used for identification of the species composition of a sample (Taberlet et al. 2012b). However, it is essential that comprehensive public databases containing well curated, up-to-date sequences from voucher specimens are available (Schenekar et al. 2020). This way, sequences generated by universal primers can be compared with the ones in reference databases, assessing sample OTU composition. The taxonomy assignment step of the eDNA metabarcoding method and thus, the identification via DNA-barcoding, is only as good and accurate as the reference databases (Cilleros et al. 2019).

Nevertheless, there is not a truly “universal” genetic marker that is capable of being amplified for all species across different taxa (Kress et al. 2015). Different markers have been used for different taxonomy groups (Deiner et al. 2017). While bacterial and archaeal diversity is often based on the 16S rRNA gene, for other taxonomic groups a diverse set of loci is used from the analogous eukaryotic rRNA gene array (e.g. ITS, 18S or 28S rRNA), chloroplast genes (for plants) and mitochondrial DNA (for eukaryotes) in an attempt for species - specific resolution (Coissac et al. 2012). The cytochrome c oxidase subunit I (COI) marker gene has been widely used for the barcoding of the Animalia kingdom for almost two decades (Hebert et al. 2003). There are cases where COI has been the standard marker for metabarcoding, such as in the assessment of freshwater macroinvertebrates (Elbrecht and Leese 2017) even though not all taxonomic groups can be differentiated to the species level using this locus (Deiner et al. 2017).

Even though there are various issues (Deagle et al. 2014), COI is indeed considered as the “gold standard” for community DNA metabarcoding of bulk metazoan samples (Andújar et al. 2018). However, as highlighted in the same study, this is not the case for eDNA samples. As Stat et al. (2017) state, in the case of eDNA samples, the target region for metazoa is found in general at considerably lower concentrations compared to those from prokaryotes because most primers targeting the COI region amplify large proportions of prokaryotes at the same time (Yang et al. 2013, 2014, Collins et al. 2019).

The co-amplification of prokaryotes is a major reason for why many Operational Taxonomic Units (OTUs) and/or Amplicon Sequence Variants (ASVs) in eDNA metabarcoding studies do not receive taxonomy assignments when metazoan reference databases are used (c.f. Aylagas et al. 2016). Despite the presence of unassigned ASVs to a varying degree in most if not all metabarcoding studies, to the best of our knowledge, there has not been a thorough investigation of the origin for these unassigned ASVs. The aim of this study was to investigate the unassigned ASVs from COI environmental data, hereafter referred to as “dark matter” as per Bernard et al. (2018), and to build a software tool for their broad taxonomic placement across the tree of life.

## Materials and methods

### Implementation

#### Building the COI tree of life

Sequences for the COI region from all the three domains of life were retrieved from curated databases. Eukaryotic sequences were retrieved from the Midori reference 2 database (version: GB239) (Machida et al. 2017). Initially, more than 1,3 millions of sequences were retrieved corresponding to 183,330 unique species from all eukaryotic taxa. With respect to bacteria and archaea, COI sequences were obtained from the BOLD database (Ratnasingham and Hebert 2007). Almost four thousand bacterial sequences were retrieved representing 559 genera. Similarly, 117 sequences from archaea were obtained from BOLD representing 41 genera. An overview of the approach that was followed is presented in Figure 1.

**Figure 1:**
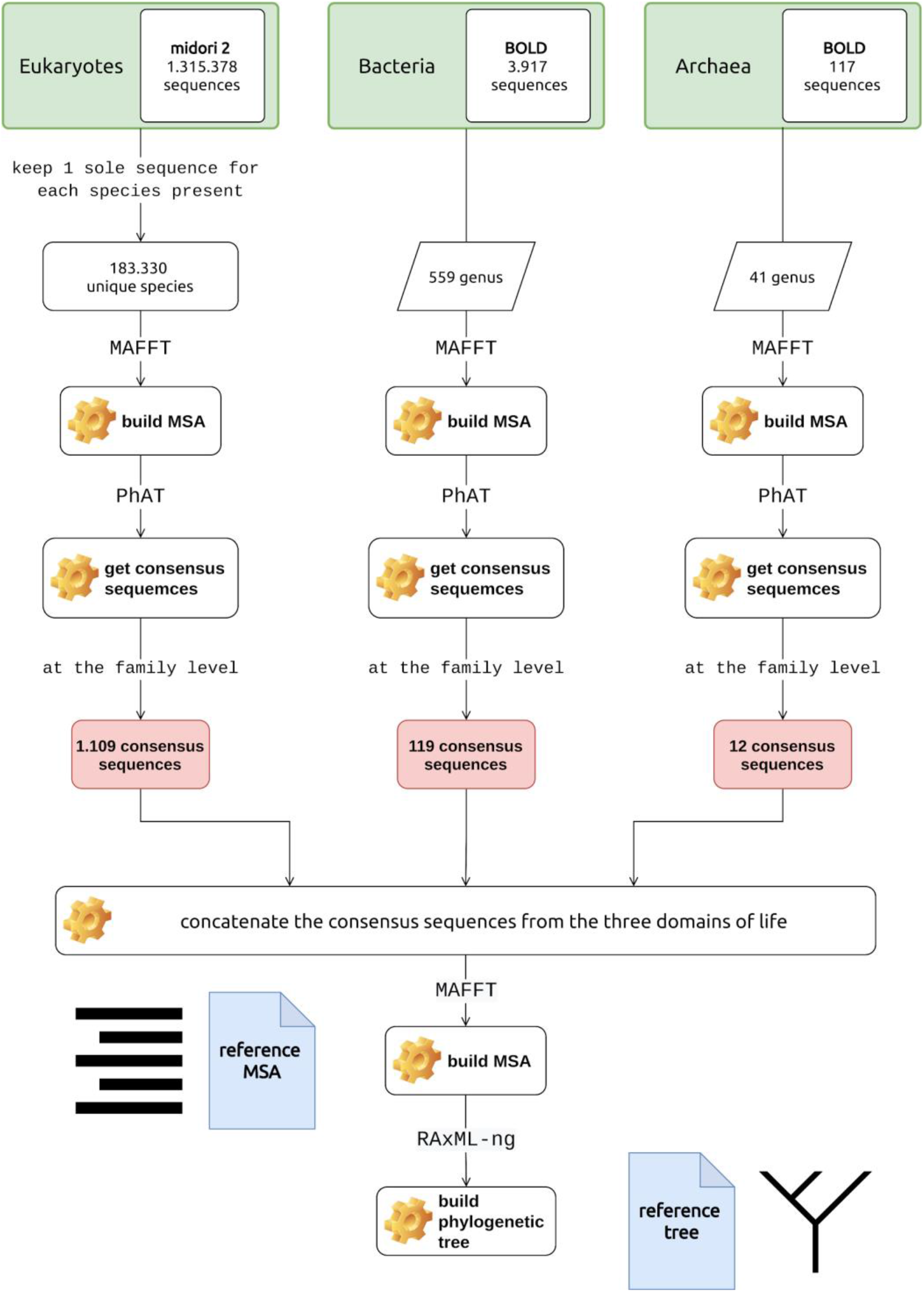
Overview of the approach followed to build the COI reference tree of life. Sequences were retrieved from: a. from Midori 2 (eukaryotes) and BOLD (bacteria and archaea) repositories. Consensus sequences at the family level were built for each domain specific dataset. `MAFFT` and consensus sequences at the family level were built using the `PhAT` algorithm. The COI reference tree was finally built using `RAxML-ng`. [Noun project](https://thenounproject.com/) icons by: by Arthur Slain and A. Beale.

The large number of obtained sequences effectively prevents a phylogenetic tree construction encompassing all of the three kingdoms (archaea, bacteria, eukaryota). Therefore, representative sequences from each of the three datasets were constructed using the `PhAT` algorithm (Czech et al. 2019); based on the entropy of a set of sequences, `PhAT` groups sequences into a given target number of groups so they reflect the diversity of all the sequences in the dataset. As PhAT uses a multiple sequence alignment (MSA) as input, all three domain-specific datasets were aligned using the `MAFFT` alignment software tool (Katoh et al. 2002). In the case of eukaryotes, the alignment of the full dataset (~183K sequences) would be impractically long. Therefore, a two-step procedure was followed; a sequence subset of 500 sequences (reference set) was selected and aligned and then used as a backbone for the alignment of all the remaining eukaryotic COI sequences. All sequences were considered reliable as they were retrieved from curated databases (Midori2 and BOLD). To build the reference set, a number (*n*) of the longest sequences from each of the various phyla were chosen, proportionally to the number (*m*) of sequences of each phylum (see Supplementary Table 1). The `--min-tax-level` parameter of the algorithm was set to correspond to the family level, forcing the PhAT algorithm to build at least one consensus sequence for all families from each domain dataset. The same approach was used for all three domain datasets.

A total of 1,109 consensus sequences were built covering the eukaryotic taxa, while 119 bacterial and 12 archaeal consensus sequences were included. These sequences were then merged as a single dataset and aligned to build a reference MSA. To this end, `MAFFT` was set to return the highest quality MSA possible. The reference tree was then built based on this MSA using the `RAxML-ng` software (Kozlov et al. 2019). `RAxML-ng` was implemented using the `GTR+FO+G4m` model and asking for 10 maximum likelihood and 10 bootstrap trees.

The methodology followed is in line with the steps described on the full stack example of the EPA-ng algorithm and a thorough description of the approach implemented to build the reference tree are presented in the [Google Collab Notebook](https://colab.research.google.com/drive/1XorHsBm1uqx5TTZsH7SeVRkUA2SS8dnY?usp=sharing). All steps for building the COI reference tree were implemented at *Zorba*, the IMBBC High Performance Computing system (Zafeiropoulos et al. 2021).

#### Investigating COI dark matter

The COI reference tree was subsequently used to build and implement the Dark mAtteR iNvestigator (`DARN`) software tool. `DARN` uses a.`fasta` file as input and returns an overview of sequence assignments per domain (eukaryotes, bacteria, archaea) after placing the query sequences of the sample on the branches of the reference tree. Sequences that are not assigned to a domain are grouped as “distant”.

The focal query sequences are aligned with respect to the reference MSA using the `PaPaRa 2.0` algorithm (Berger and Stamatakis 2012). The query sequences are then split to build a discrete query MSA. Finally, the Evolutionary Placement Algorithm `EPA-ng` (Barbera et al. 2019) is used to assign the query sequences to the reference tree.

To visualise the query sequence assignments, a two-step method was developed. First, `DARN` invokes the gappa assign tool which taxonomically assigns placed query sequences by making use of the likelihood weight (LWR) that was assigned to this exact taxonomic path. In the `DARN` framework, by making use of the `--per-query-results` and `--best-hit` flags, the `gappa assign` software assigns the LWR of each placement of the query sequences to a taxonomic rank that was built based on the taxonomies included in the reference tree. The first flag ensures that the `gappa assign` tool will return a tabular file containing one assignment profile per input query while the latter will only return the assignment with the highest LWR. `DARN` automatically parses this output of `gappa assign` to build two input Krona (Ondov et al. 2011) profile files based on a) the LWR values of each query sequence and b) an adjustive approach where all the best hits get the same value in a binary approach (presence - absence). In the `final_outcome` directory that DARN creates, `two .html` files, one for each of the Krona plots. In addition four .`fasta` files are generated including the sequences of the sample that have been assigned to each domain or as “distant”. A .`json` file with the metadata of the analysis is also returned including the identities of the sequences assigned to each domain.

`DARN` also runs the `gappa assign` tool with the `--per-query-results` flag only. This way, the user can have a thorough overview of each sample’s sequence assignments, as a sequence may be assigned to more than one branch of the reference tree, sometimes even to different domains.

`DARN` source code is available on [GitHub](https://github.com/hariszaf/darn/tree/main).

### Software evaluation

To evaluate DARN on the presence of dark matter we analysed a wide range of cases to show the ability of DARN to detect and estimate dark matter under various conditions. Both eDNA and bulk samples, from marine, lotic and lentic environments, were selected to reflect various combinations of primer and amplicon lengths, PCR protocols and bioinformatics analyses (Table 1).

More specifically, 57 marine, surface water, eDNA samples from Ireland were analysed through a. QIIME2 (Bolyen et al. 2018) and DADA2 (Callahan et al. 2016) and, b. PEMA (Zafeiropoulos et al. 2020). Similarly, 18 mangrove and 18 reef marine eDNA samples from Honduras, were analyzed using a. JAMP v0.74 (Elbrecht 2021) and DnoisE (Antich et al. 2021) and b. PEMA . Furthermore, a sediment sample and two samples from Autonomous Reef Monitoring Structures (ARMS) one conserved in DMSO and another in ethanol from the Obst et al. (2020) dataset were analysed using PEMA. In addition, one lotic and two lentic samples from Norway were analysed using PEMA. For the case of the lentic samples, multiple parameter sets regarding the ASVs inference step were implemented; i.e the *d* parameter of the Swarm v2 (Mahé et al. 2015) that PEMA invokes was set equal to 2 and 10 to cover a great range of different cases (Kamenova 2020).

DARN was then executed using the ASVs retrieved in each case as input. All the DARN analyses and the PEMA runs were performed on an Intel(R) Xeon(R) CPU E5649 @ 2.53GHz server of 24 CPUs and 142 GB RAM in the Area52 Research Group at the University College Dublin.

## Results and Discussion

Most, if not all, eDNA metabarcoding approaches will amplify the target taxa but they will also often result in a significantly large proportion of unassignable sequences with unknown origin, i.e. dark matter. The dark matter is rarely further analysed but has been suspected of containing bacteria, algae and fungi (Yang et al. 2013, Aylagas et al. 2016). Although the unassignable sequences could be informative, there have been few attempts to further investigate the dark matter (e.g. (Sinniger et al. 2016, Haenel et al. 2017)). In an effort to shine light on the contents of dark matter we present a bioinformatics pipeline - DARN, that is capable of assigning sequences from COI based metabarcoding to taxa representing the entire tree of life. A Docker image is now available to share DARN as a one-stop-shop tool for the quantification and the investigation of dark matter in sequence samples. It is important to note that DARN was not designed to be a taxonomy assignment method, but to provide a generic view of the OTUs/ASVs retrieved after the bioinformatics analyses of environmental samples. The approach allows, once the bioinformatics analyses are completed, for sequences with no taxonomic assignments to be further investigated.

A COI-oriented reference phylogenetic tree of life was built by using 1,240 consensus sequences with more than 80% of those coming from eukaryotic taxa. The number of sequences returned after the bioinformatics analysis of a series of samples using the bioinformatic analyses ranged from c. 3k to 214k (Table 2) workflow. A coherent visual representation of the DARN outcome for all the datasets is available at https://hariszaf.github.io/darn/.

**Table 2:**
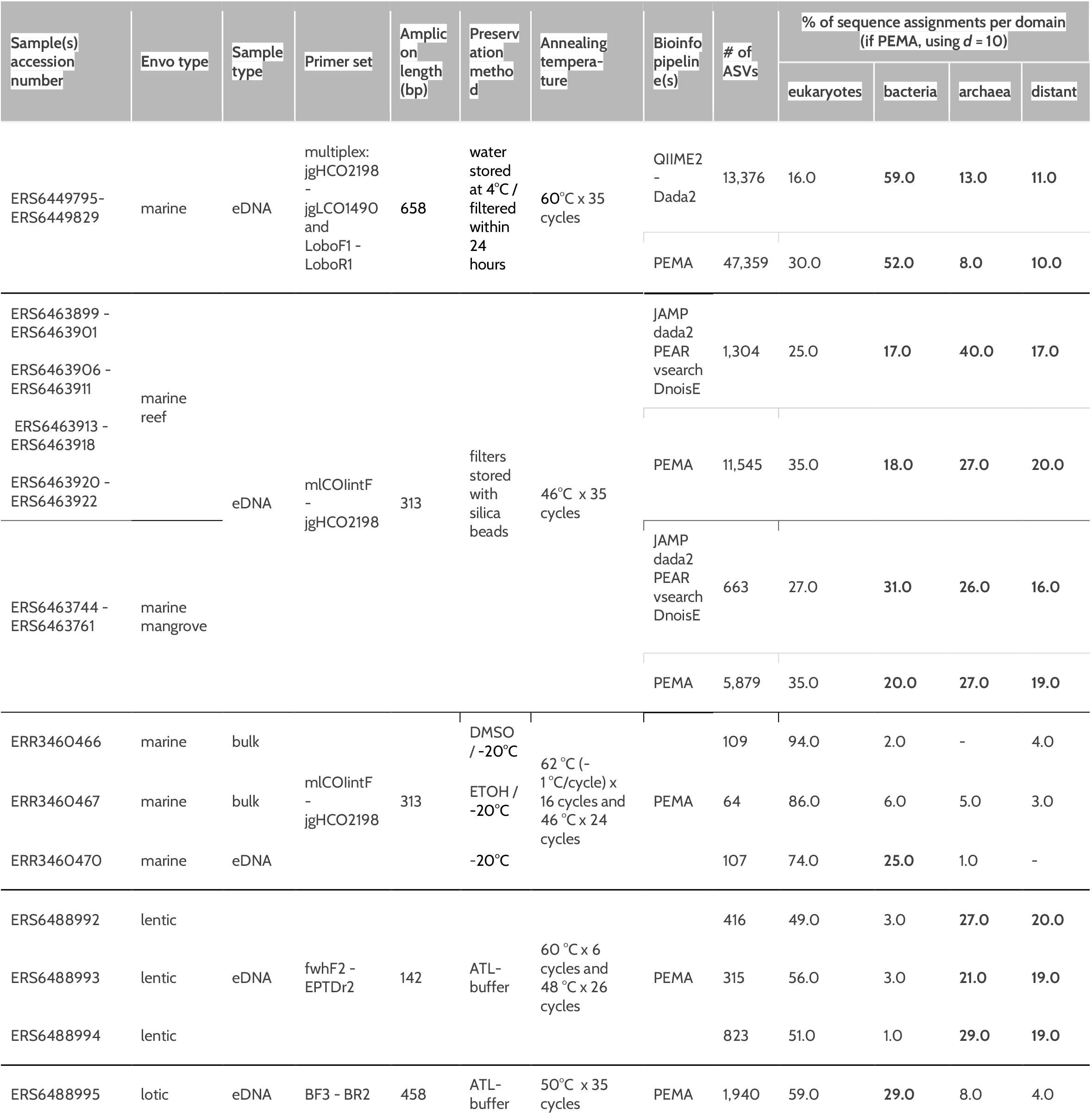
DARN outcome over the samples or set of samples. Assignment fractions of the sequences per domain per sample in the DARN results over the samples.

Our results clearly demonstrate that standard metabarcoding approaches based on the COI gene region of the mitochondrial genome will not only amplify eukaryotes, but also a large proportion of non-target prokaryotic organisms, such as bacteria and archaea.

Significant proportions of non-eukaryote DARN assignments were observed in all marine eDNA samples (Table 2). Bacterial assignments made up the largest proportion of the non-eukaryotic assignments (~50% of the ASVs in some cases), however, archaeal assignments were also detected to a great extent as well (~20% on average). The lentic samples were those with the shortest amplicon length among those analysed (142bp). In all the three samples analysed, the percentage of eukaryote assignments were around 50%; bacteria, archaea and “distant” assignments had similar percentages too. In the case of bulk samples (Table 2) only a low proportion of the sequences were not assigned as eukaryotes, suggesting that non-eukaryotic sequences are more abundant in environmental samples. This could be expected since prokaryotes are amplified as whole organisms from environmental samples, while eukaryotes are amplified from their DNA traces and not from any obvious biological source material.

While our approach specifically addressed the COI gene, DARN can be adapted to analyse any locus fragment. For instance, metabarcoding of environmental samples for the 12S rRNA mitochondrial region is often employed to assess fish biodiversity (Weigand et al. 2019, Miya et al. 2020) and the approach presented here could be adjusted to allow further analyses of the 12S rRNA data.

As publicly available bacterial COI sequences are far too few to represent the bacterial variability, their reliable taxonomic identification is not possible currently. Therefore, we suggest bacterial COI sequences to be included in the reference databases used for the taxonomy assignment of OTUs/ASVs in COI-based eukaryote metabarcoding studies. This way, bacterial non-target sequences that were amplified during the library preparation can have a taxonomy assignment.

## Conclusions

This study presents a software tool (DARN) to enable the investigation of the unknown, unassigned COI sequences (OTUs/ASVs) derived from eDNA and bulk DNA metabarcoding studies. DARN enables the classification of the unassigned sequences to the three domains of life or in a distinct group called “distant” when a sequence phylogenetically differs to a great extent from all of them. The Krona plots that DARN builds provide a graphical and numeric overview of the sample, showing the assignments to the various groups, while the tabular files returned allow for further analyses of the assigned sequences. Clearly, dark matter, and especially bacteria, make up a significant proportion of sequences generated in COI based eDNA metabarcoding datasets. The large proportion of prokaryotes observed in the present study is corroborated by the findings of Yang et al. (2013). Furthermore, dark matter seems to be particularly common in eDNA as compared to bulk samples (Andújar et al. 2018). Our implementations using DARN indicate that it is essential both for the global reference databases (e.g BOLD, Midori etc) and custom reference databases which are commonly used, to also include non-eukaryotic sequences. The visual and interactive properties of the Krona plot allows the user to navigate through the taxonomy and also extract fasta sequences headers from the different assigned taxa for further analyses. Furthermore, DARN also supports a thorough investigation per OTU/ASV, as it returns a .json file with all the OTUs/ASVs ids that have been assigned in each of the four categories (bacteria, archaea, eukaryotes and distant).

The approaches implemented in DARN can benefit both bulk and eDNA metabarcoding studies, by allowing quality control and further investigation of the unassigned OTUs/ASVs. The approach is also adaptable to other markers than COI. Moreover, the approach presented here allows researchers to better understand the unknown unknowns and shed light on the dark matter of their metabarcoding sequence data.

### Licence

License: GNU GPLv3. For third-party components separate licenses apply.

### CRediT

H.Z. → Conceptualization, Methodology, Software, Validation, Investigation, Resources, Writing - Original draft

L.G. → Methodology, Validation, Investigation

S.H. → Validation, Investigation

C.P. → Validation, Investigation, Writing - Original draft

J.C. → Conceptualization, Investigation, Resources, Writing - Original draft

## Availability

GitHub repo: https://github.com/hariszaf/darn

DockerHub repo: https://hub.docker.com/r/hariszaf/darn

The sequence data that support the findings of this study are available in the European Nucleotide Archive (ENA) with the following study accession numbers:

- Marine samples from Ireland: PRJEB45030
- Marine samples from Honduras: PRJEB45038
- Marine ARMS samples: PRJEB33796
- Lake and riverine samples from Norway: PRJEB45246

## Acknowledgement

This research was supported in part through computational resources provided by IMBBC (Institute of Marine Biology, Biotechnology and Aquaculture) of the HCMR (Hellenic Centre for Marine Research). Funding for establishing the IMBBC HPC has been received by the MARBIGEN (EU Regpot) project, LifeWatchGreece RI and the CMBR (Centre for the study and sustainable exploitation of Marine Biological Resources) RI.

We would also like to thank Dr. Evangelos Pafilis for providing us access to the IMBBC HPC infrastructure and Dr. Frode Fossøy for providing us environmental, lake and riverine sequence samples from Norway.

## Supplementary Files

**Supplementary Table 1:** Number of sequences per phylum in the eukaryotes domain dataset and the corresponding number of the longest sequences used in the 500 sequences subset (reference set) used as a backbone for the complete alignment.

## References

Andruszkiewicz EA, Starks HA, Chavez FP, Sassoubre LM, Block BA, Boehm AB (2017) Biomonitoring of marine vertebrates in Monterey Bay using eDNA metabarcoding. PLOS ONE 12: e0176343. https://doi.org/10.1371/journal.pone.0176343

Andújar C, Arribas P, Yu DW, Vogler AP, Emerson BC (2018) Why the COI barcode should be the community DNA metabarcode for the metazoa. Molecular Ecology 27: 3968–3975. https://doi.org/10.1111/mec.14844

Antich A, Palacin C, Wangensteen OS, Turon X (2021) To denoise or to cluster, that is not the question: optimizing pipelines for COI metabarcoding and metaphylogeography. BMC Bioinformatics 22: 177. https://doi.org/10.1186/s12859-021-04115-6

Aylagas E, Borja Á, Irigoien X, Rodríguez-Ezpeleta N (2016) Benchmarking DNA Metabarcoding for Biodiversity-Based Monitoring and Assessment. Frontiers in Marine Science 3. https://doi.org/10.3389/fmars.2016.00096

Barbera P, Kozlov AM, Czech L, Morel B, Darriba D, Flouri T, Stamatakis A (2019) EPA-ng: Massively Parallel Evolutionary Placement of Genetic Sequences. Systematic Biology 68: 365–369. https://doi.org/10.1093/sysbio/syy054

Berger SA, Stamatakis A (2012) PaPaRa 2.0: A Vectorized Algorithm for Probabilistic Phylogeny-Aware Alignment Extension. Heidelberg Institute for Theoretical Studies: 12.

Bernard G, Pathmanathan JS, Lannes R, Lopez P, Bapteste E (2018) Microbial Dark Matter Investigations: How Microbial Studies Transform Biological Knowledge and Empirically Sketch a Logic of Scientific Discovery. Genome Biology and Evolution 10: 707–715. https://doi.org/10.1093/gbe/evy031

Bolyen E, Rideout JR, Dillon MR, Bokulich NA, Abnet C, Al-Ghalith GA, Alexander H, Alm EJ, Arumugam M, Asnicar F, Bai Y, Bisanz JE, Bittinger K, Brejnrod A, Brislawn CJ, Brown CT, Callahan BJ, Caraballo-Rodríguez AM, Chase J, Cope E, Silva RD, Dorrestein PC, Douglas GM, Durall DM, Duvallet C, Edwardson CF, Ernst M, Estaki M, Fouquier J, Gauglitz JM, Gibson DL, Gonzalez A, Gorlick K, Guo J, Hillmann B, Holmes S, Holste H, Huttenhower C, Huttley G, Janssen S, Jarmusch AK, Jiang L, Kaehler B, Kang KB, Keefe CR, Keim P, Kelley ST, Knights D, Koester I, Kosciolek T, Kreps J, Langille MG, Lee J, Ley R, Liu Y-X, Loftfield E, Lozupone C, Maher M, Marotz C, Martin BD, McDonald D, McIver LJ, Melnik AV, Metcalf JL, Morgan SC, Morton J, Naimey AT, Navas-Molina JA, Nothias LF, Orchanian SB, Pearson T, Peoples SL, Petras D, Preuss ML, Pruesse E, Rasmussen LB, Rivers A, Michael S Robeson II, Rosenthal P, Segata N, Shaffer M, Shiffer A, Sinha R, Song SJ, Spear JR, Swafford AD, Thompson LR, Torres PJ, Trinh P, Tripathi A, Turnbaugh PJ, Ul-Hasan S, Hooft JJ van der, Vargas F, Vázquez-Baeza Y, Vogtmann E, Hippel M von, Walters W, Wan Y, Wang M, Warren J, Weber KC, Williamson CH, Willis AD, Xu ZZ, Zaneveld JR, Zhang Y, Zhu Q, Knight R, Caporaso JG (2018) QIIME 2: Reproducible, interactive, scalable, and extensible microbiome data science. PeerJ Inc. https://doi.org/10.7287/peerj.preprints.27295v2

Callahan BJ, McMurdie PJ, Rosen MJ, Han AW, Johnson AJA, Holmes SP (2016) DADA2: High-resolution sample inference from Illumina amplicon data. Nature Methods 13: 581–583. https://doi.org/10.1038/nmeth.3869

Cilleros K, Valentini A, Allard L, Dejean T, Etienne R, Grenouillet G, Iribar A, Taberlet P, Vigouroux R, Brosse S (2019) Unlocking biodiversity and conservation studies in high-diversity environments using environmental DNA (eDNA): A test with Guianese freshwater fishes. Molecular Ecology Resources 19: 27–46. https://doi.org/10.1111/1755-0998.12900

Coissac E, Riaz T, Puillandre N (2012) Bioinformatic challenges for DNA metabarcoding of plants and animals. Molecular Ecology 21: 1834–1847. https://doi.org/10.1111/j.1365-294X.2012.05550.x

Collins RA, Bakker J, Wangensteen OS, Soto AZ, Corrigan L, Sims DW, Genner MJ, Mariani S (2019) Non-specific amplification compromises environmental DNA metabarcoding with COI. Methods in Ecology and Evolution 10: 1985–2001. https://doi.org/10.1111/2041-210X.13276

Czech L, Barbera P, Stamatakis A (2019) Methods for automatic reference trees and multilevel phylogenetic placement. Bioinformatics 35: 1151–1158. https://doi.org/10.1093/bioinformatics/bty767

Deagle BE, Jarman SN, Coissac E, Pompanon F, Taberlet P (2014) DNA metabarcoding and the cytochrome c oxidase subunit I marker: not a perfect match. Biology Letters 10: 20140562. https://doi.org/10.1098/rsbl.2014.0562

Deiner K, Bik HM, Mächler E, Seymour M, Lacoursière‐Roussel A, Altermatt F, Creer S, Bista I, Lodge DM, Vere N de, Pfrender ME, Bernatchez L (2017) Environmental DNA metabarcoding: Transforming how we survey animal and plant communities. Molecular Ecology 26: 5872–5895. https://doi.org/10.1111/mec.14350

Elbrecht V (2021) VascoElbrecht/JAMP. . R. Available from: https://github.com/VascoElbrecht/JAMP (May 28, 2021).

Elbrecht V, Leese F (2017) Validation and Development of COI Metabarcoding Primers for Freshwater Macroinvertebrate Bioassessment. Frontiers in Environmental Science 5. https://doi.org/10.3389/fenvs.2017.00011

Haenel Q, Holovachov O, Jondelius U, Sundberg P, Bourlat SJ (2017) NGS-based biodiversity and community structure analysis of meiofaunal eukaryotes in shell sand from Hållö island, Smögen, and soft mud from Gullmarn Fjord, Sweden. Biodiversity Data Journal. https://doi.org/10.3897/BDJ.5.e12731

Hebert PDN, Ratnasingham S, Waard JR de (2003) Barcoding animal life: cytochrome c oxidase subunit 1 divergences among closely related species. Proceedings of the Royal Society of London. Series B: Biological Sciences. https://doi.org/10.1098/rsbl.2003.0025

Ji Y, Ashton L, Pedley SM, Edwards DP, Tang Y, Nakamura A, Kitching R, Dolman PM, Woodcock P, Edwards FA, Larsen TH, Hsu WW, Benedick S, Hamer KC, Wilcove DS, Bruce C, Wang X, Levi T, Lott M, Emerson BC, Yu DW (2013) Reliable, verifiable and efficient monitoring of biodiversity via metabarcoding. Ecology Letters 16: 1245–1257. https://doi.org/10.1111/ele.12162

Kamenova S (2020) A flexible pipeline combining clustering and correction tools for prokaryotic and eukaryotic metabarcoding. Peer Community in Ecology 1: 100043. https://doi.org/10.24072/pci.ecology.100043

Katoh K, Misawa K, Kuma K, Miyata T (2002) MAFFT: a novel method for rapid multiple sequence alignment based on fast Fourier transform. Nucleic Acids Research 30: 3059–3066. https://doi.org/10.1093/nar/gkf436

Kozlov AM, Darriba D, Flouri T, Morel B, Stamatakis A (2019) RAxML-NG: A fast, scalable, and user-friendly tool for maximum likelihood phylogenetic inference. bioRxiv: 447110. https://doi.org/10.1101/447110

Machida RJ, Leray M, Ho S-L, Knowlton N (2017) Metazoan mitochondrial gene sequence reference datasets for taxonomic assignment of environmental samples. Scientific Data 4: 1–7. https://doi.org/10.1038/sdata.2017.27

Mahé F, Rognes T, Quince C, Vargas C de, Dunthorn M (2015) Swarm v2: highly-scalable and high-resolution amplicon clustering. PeerJ 3: e1420. https://doi.org/10.7717/peerj.1420

Miya M, Gotoh RO, Sado T (2020) MiFish metabarcoding: a high-throughput approach for simultaneous detection of multiple fish species from environmental DNA and other samples. Fisheries Science 86: 939–970. https://doi.org/10.1007/s12562-020-01461-x

Obst M, Exter K, Allcock AL, Arvanitidis C, Axberg A, Bustamante M, Cancio I, Carreira-Flores D, Chatzinikolaou E, Chatzigeorgiou G, Chrismas N, Clark MS, Comtet T, Dailianis T, Davies N, Deneudt K, de Cerio OD, Fortič A, Gerovasileiou V, Hablützel PI, Keklikoglou K, Kotoulas G, Lasota R, Leite BR, Loisel S, Lévêque L, Levy L, Malachowicz M, Mavrič B, Meyer C, Mortelmans J, Norkko J, Pade N, Power AM, Ramšak A, Reiss H, Solbakken J, Staehr PA, Sundberg P, Thyrring J, Troncoso JS, Viard F, Wenne R, Yperifanou EI, Zbawicka M, Pavloudi C (2020) A Marine Biodiversity Observation Network for Genetic Monitoring of Hard-Bottom Communities (ARMS-MBON). Frontiers in Marine Science 7. https://doi.org/10.3389/fmars.2020.572680

Ondov BD, Bergman NH, Phillippy AM (2011) Interactive metagenomic visualization in a Web browser. BMC Bioinformatics 12: 385. https://doi.org/10.1186/1471-2105-12-385

Ratnasingham S, Hebert PDN (2007) bold: The Barcode of Life Data System (http://www.barcodinglife.org). Molecular Ecology Notes 7: 355–364. https://doi.org/10.1111/j.1471-8286.2007.01678.x

Ruppert KM, Kline RJ, Rahman MS (2019) Past, present, and future perspectives of environmental DNA (eDNA) metabarcoding: A systematic review in methods, monitoring, and applications of global eDNA. Global Ecology and Conservation 17: e00547. https://doi.org/10.1016/j.gecco.2019.e00547

Schenekar T, Schletterer M, Lecaudey LA, Weiss SJ (2020) Reference databases, primer choice, and assay sensitivity for environmental metabarcoding: Lessons learnt from a re-evaluation of an eDNA fish assessment in the Volga headwaters. River Research and Applications 36: 1004–1013. https://doi.org/10.1002/rra.3610

Sinniger F, Pawlowski J, Harii S, Gooday AJ, Yamamoto H, Chevaldonné P, Cedhagen T, Carvalho G, Creer S (2016) Worldwide Analysis of Sedimentary DNA Reveals Major Gaps in Taxonomic Knowledge of Deep-Sea Benthos. Frontiers in Marine Science 3. https://doi.org/10.3389/fmars.2016.00092

Stat M, Huggett MJ, Bernasconi R, DiBattista JD, Berry TE, Newman SJ, Harvey ES, Bunce M (2017) Ecosystem biomonitoring with eDNA: metabarcoding across the tree of life in a tropical marine environment. Scientific Reports 7: 12240. https://doi.org/10.1038/s41598-017-12501-5

Taberlet P, Coissac E, Hajibabaei M, Rieseberg LH (2012a) Environmental DNA. Molecular Ecology 21: 1789–1793. https://doi.org/10.1111/j.1365-294X.2012.05542.x

Taberlet P, Coissac E, Pompanon F, Brochmann C, Willerslev E (2012b) Towards next-generation biodiversity assessment using DNA metabarcoding. Molecular Ecology 21: 2045–2050. https://doi.org/10.1111/j.1365-294X.2012.05470.x

Weigand H, Beermann AJ, Čiampor F, Costa FO, Csabai Z, Duarte S, Geiger MF, Grabowski M, Rimet F, Rulik B, Strand M, Szucsich N, Weigand AM, Willassen E, Wyler SA, Bouchez A, Borja A, Čiamporová-Zaťovičová Z, Ferreira S, Dijkstra K-DB, Eisendle U, Freyhof J, Gadawski P, Graf W, Haegerbaeumer A, van der Hoorn BB, Japoshvili B, Keresztes L, Keskin E, Leese F, Macher JN, Mamos T, Paz G, Pešić V, Pfannkuchen DM, Pfannkuchen MA, Price BW, Rinkevich B, Teixeira MAL, Várbíró G, Ekrem T (2019) DNA barcode reference libraries for the monitoring of aquatic biota in Europe: Gap-analysis and recommendations for future work. Science of The Total Environment 678: 499–524. https://doi.org/10.1016/j.scitotenv.2019.04.247

Yang C, Ji Y, Wang X, Yang C, Yu DW (2013) Testing three pipelines for 18S rDNA-based metabarcoding of soil faunal diversity. Science China Life Sciences 56: 73–81. https://doi.org/10.1007/s11427-012-4423-7

Yang C, Wang X, Miller JA, de Blécourt M, Ji Y, Yang C, Harrison RD, Yu DW (2014) Using metabarcoding to ask if easily collected soil and leaf-litter samples can be used as a general biodiversity indicator. Ecological Indicators 46: 379–389. https://doi.org/10.1016/j.ecolind.2014.06.028

Zafeiropoulos H, Viet HQ, Vasileiadou K, Potirakis A, Arvanitidis C, Topalis P, Pavloudi C, Pafilis E (2020) PEMA: a flexible Pipeline for Environmental DNA Metabarcoding Analysis of the 16S/18S ribosomal RNA, ITS, and COI marker genes. GigaScience 9. https://doi.org/10.1093/gigascience/giaa022

Zafeiropoulos H, Gioti A, Ninidakis S, Potirakis A, Paragkamian S, Angelova N, Antoniou A, Danis T, Kaitetzidou E, Kasapidis P, Kristoffersen JB, Papadogiannis V, Pavloudi C, Ha QV, Lagnel J, Pattakos N, Perantinos G, Sidirokastritis D, Vavilis P, Kotoulas G, Manousaki T, Sarropoulou E, Tsigenopoulos CS, Arvanitidis C, Magoulas A, Pafilis E (2021) The IMBBC HPC facility: history, configuration, usage statistics and related activities. Zenodo https://doi.org/10.5281/zenodo.4665308

